# A Distinct Subpopulation of Extended Amygdala Neurons Drives Food Intake

**DOI:** 10.1101/2025.11.07.685941

**Authors:** Isaac F. Kandil, Ethan T. Rogers, Allison R. Morningstar, William J. Giardino

## Abstract

**Background:** Neurons in the oval subnucleus of the bed nucleus of the stria terminalis (ovBNST) integrate stress and reward signals to regulate motivated behaviors, including food consumption. However, the contribution of specific ovBNST neuronal subpopulations remains poorly understood. Here, we investigated vasoactive intestinal peptide receptor 2 *(Vipr2)* expressing ovBNST neurons using chemogenetics, immunohistochemistry, and viral circuit mapping.

**Methods and Results:** Using stimulatory hM3Dq designer receptors exclusively activated by designer drugs (DREADDs), we found that chemogenetic activation of ovBNST*^Vipr2^*neurons significantly increased food intake. We then quantified cFos activation in *Vipr2*-tdTomato reporter mice following several unique feeding-related manipulations, finding that food restriction (FR) robustly activated ovBNST*^Vipr2^* neurons. Further analysis revealed decreased vasoactive intestinal peptide (VIP) innervation of the ovBNST following FR, in which reduced VIP expression was significantly associated with greater ovBNST*^Vipr2^* cFos activation. Given previous reports of reduced food intake following stimulation of ovBNST neurons expressing protein kinase C delta (PKCδ), we used immunostaining to uncover that *Vipr2* and PKCδ mark largely non-overlapping ovBNST neuronal subpopulations, aligning with their opposing effects on food intake. Finally, Cre-dependent anterograde viral tracing revealed that ovBNST*^Vipr2^* neurons project prominently to the parasubthalamic nucleus (PSTN) and paraventricular nucleus of the hypothalamus (PVN), two feeding-related regions.

**Conclusions:** Together, these results identify ovBNST*^Vipr2^*neurons as a functionally distinct BNST subpopulation that promotes feeding, is activated by food restriction, and links ovBNST neuropeptide signaling to hypothalamic feeding centers.

## INTRODUCTION

Several neural systems are involved in the regulation of appetite and food consumption. The bed nucleus of the stria terminalis (BNST) is most known for regulating anxiety and stress but has also been implicated in other behaviors relevant to psychiatric disorders, including feeding and binge intake of reinforcing substances [1, 2]. In particular, excitotoxic lesion of the oval subnucleus of the BNST (ovBNST) attenuated stress-induced weight gain in rats [3], stimulation of BNST-Vglut3 neurons reduced sucrose intake in fasted mice [4], and stimulation of GABAergic BNST neurons projecting to the lateral hypothalamus (LH) or anterior periaqueductal gray rapidly promoted feeding [5–8]. However, the BNST exhibits remarkable heterogeneity in regard to its neuronal sub-types and projection patterns. For example, stimulation of ovBNST neurons containing protein kinase C delta (PKCd) reduce food intake [9, 10], in opposition to the larger GABAergic population. Overall, the precise contribution of the BNST to different feeding-related phenotypes is highly dependent on which neurotransmitters and neuropeptides are expressed by the individual cell population, including heterogeneity within the ovBNST itself [11–14].

Regarding ovBNST heterogeneity with potential contributions to food intake, transcriptomic data indicates that *Vipr2*, the gene encoding the vasoactive intestinal peptide receptor 2, is a marker for a distinct subset of ovBNST neurons [15], and the VIP system has been broadly implicated in appetite regulation [16]. In fact, VIP and Vipr2 are part of a larger neuropeptide system including pituitary adenylyl cyclase-activating polypeptide (PACAP), PACAP type-1 receptor (Pacr1), and Vipr1. Pacr1 can be bound by PACAP selectively, while Vipr1 and Vipr2 can be bound by either VIP or PACAP. PACAP/Pacr1 signaling in the BNST generally promotes anxiety-related stress responses [17–19] and alcohol drinking [20], while VIP/Vipr2 BNST signaling is less well studied. Although previous research identified that VIP innervation of the ovBNST was reduced in obese mice chronically maintained on high-fat diet [21], no investigations have directly tested the feeding-related functions of VIP and Vipr2 signaling in the ovBNST.

Here, we obtained genetic access to Vipr2 neurons in the ovBNST using *Vipr2*-Cre mice and performed chemogenetic stimulation of ovBNST*^Vipr2^*neurons while measuring food intake. We assessed neuronal activity in ovBNST*^Vipr2^* neurons with cFos immunostaining following exposure to various feeding-related stimuli, characterized overlap of ovBNST*^Vipr2^*neurons with a marker for known appetite-suppressing ovBNST neurons, and visualized ovBNST*^Vipr2^* neuron axonal outputs to downstream nuclei. Overall, our findings support a framework in which ovBNST*^Vipr2^*neurons promote feeding (in contrast to non-overlapping ovBNST PKCd neurons), are activated by food restriction (in proportion to axonal VIP innervation from an as-yet-unknown source), and project to canonical feeding-related structures in the hypothalamus.

## METHODS

### Animals

All experiments were performed with approval by Stanford University’s Administrative Panel on Laboratory Animal Care (APLAC #33087). All experimental subjects that underwent surgery for behavioral and circuit tracing experiments were *Vipr2*^em1.1(cre)Hze^ mice (Strain #031332) generated on a C57BL/6J background and at least 8 weeks old at the time of surgeries. *Vipr2*-Cre mice were crossed with B6.Cg-Gt(ROSA)26Sor^tm14(CAG-tdTomato)Hze^ mice to produce *Vipr2*::Ai14 that selectively express tdTomato in neurons expressing Cre driven by *Vipr2*.

### Surgeries

Mice underwent bilateral stereotaxic viral injection surgeries for chemogenetic/anterograde tracing experiments as described [22–26]. Mice were anesthetized with ketamine and xylazine (100 mg/kg ketamine; 20 mg/kg xylazine, i.p.) before being placed on a stereotaxic frame. Viruses were bilaterally injected at the BNST (+0.35 A/P, +/-M/L 0.89, D/V –4.22 mm from bregma) at a rate of 75nl/min with a 28G cannula (Component supply; HTXC1-28R). The cannula was kept in place for 5 minutes before being slowly retracted to allow for viral infusion. All adeno-associated viruses (AAVs) were obtained from the Gene Vector and Virus Core at Stanford University. *Vipr2*-Cre mice that underwent DREADD manipulations received bilateral injections (250 nl/hemisphere) of AAVdj-EF1a-DIO-hM3D(Gq)-mCherry (AAV-130, 1 x 10^12^ vg/ml) or AAVdj-EF1a-DIO-mCherry (AAV-14, 3.2 x 10^12^ vg/ml). Anterograde tracing experiments used AAV-DJ-hSyn-FLEX-mGFP-2A Synaptophysin-mRuby (lot #6774, 1.6 x 10^13^ vg/ml, 250 nl/hemisphere).

### Behavioral Procedures – Food intake

To reversibly activate ovBNST*^Vipr2^* neurons, we performed chemogenetic manipulations using DREADDs. Mice received bilateral injections of DREADDs in the BNST to selectively express Gq-coupled human M3 muscarinic receptor (hM3Dq), an excitatory DREADD, in ovBNST*^Vipr2^* neurons. Mice that were injected with hM3Dq were given two weeks to recover from surgery before drinking experiments. Mice underwent a standard Drinking in the Dark (DID) 2-bottle-choice binge alcohol drinking paradigm as described [27, 28]. During week 1, no alcohol (EtOH) was given to obtain a water baseline with two identical bottles. Water and food intakes were recorded on four consecutive days in a four-hour period from ZT14-ZT18, an interval of the circadian cycle when fluid consumption levels are near maximal. Three days after the end of week 1, week 2 began with an identical protocol except one bottle was replaced with 20% EtOH during the 4hr drinking sessions, and we continued to measure food intake during the daily 4hr sessions. 4hr ethanol consumption (ml) was converted to grams based on density of EtOH concentration and divided by the animal’s body weight to give daily ethanol intake expressed as g/kg. 4hr water, total liquid (ethanol + water), and food consumption were divided by the animal’s body weight to give 4hr water and total liquid intake expressed as ml/kg and 4hr food intake expressed as g/kg. 4hr ethanol preference was calculated by dividing total ethanol consumption (ml) by total liquid consumption (ml) and converting to a percentage.

### Immunostaining

For cFos staining in Gq-DREADD and virus groups, mice received i.p. Injections of saline or DCZ (0.1 mg/kg) 90mins prior to perfusion. For perfusion, mice were anesthetized with ketamine and xylazine (100 mg/kg ketamine; 20 mg/kg xylazine, i.p.) and perfused transcardially with 1x phosphate buffered saline (PBS), followed by 4% paraformaldehyde (PFA) in PBS. Brains were extracted and fixed overnight (12-18h) in a 4% PFA in PBS solution at 4°C, and then were placed in a sucrose solution (30% sucrose in PBS with 0.1% NaN3) for 48-96h at 4°C. Brains were then sliced in 30 μm coronal sections on a microtome (Epredia; HM 450 Sliding Microtome to be collected in 24-well plates containing a 0.1% Sodium Azide (NaN3) solution in PBS. Well plates were covered and stored at 4°C until imaging and/or immunohistochemical processing as described [24, 25].

Slices were washed in PBS for 5 minutes before being incubated for 1 hour in a blocking solution of 0.3% Triton X-100 in 1% PSB (PBST) containing 4% bovine serum albumin (BSA). Slices were then incubated with the primary antibody overnight (12-18hrs) in 4% BDA/PBST blocking solution. The slices were washed in three steps in PBS before being incubated for 1 hour before adding the secondary antibodies and incubating for an additional 2+ hours. Slices were washed again 3x in PBS and were then mounted onto gelatin-coated slides (Lab Scientific; 7802) and coverslipped with Fluoroshield containing DAPI Mounting Media (Sigma; F6057).

To validate Gq-DREADD activation in *Vipr2*-Cre mice and feeding-related stimuli neuronal activation in *Vipr2*-tdTomato mice, we performed immunohistochemical staining for the immediately early gene c-Fos that marks strong neuronal activation. We used rabbit anti-cFos primary antibody (EnCore Biotechnology, catalog # RPCA-c-FOS, 1:2500) and donkey anti-rabbit 488 secondary (Abcam #ab150073, 1:500). To compare PKCd and *Vipr2*-tdTomato expression, we used mouse anti-PKCd (BD Transduction #610397, 1:500) and goat anti-mouse 488 secondary (ThermoFisher #A-11001, 1:500).

All images were obtained on a Zeiss LSM 900 Confocal Microscope using ZEN software. For anterograde tracing analysis, all images were analyzed using FiJi to quantify fluorescence.

### Feeding-Related Stimuli for Vipr2-tomato cFos Staining

For Food Restriction (FR) stimuli, all food was removed from cages 24hours before perfusion. For High Fat Diet (HFD) stimulus, mice were exposed to HFD pellets (Inotiv, TD.88137) for 3 consecutive days prior to perfusion through a perforated container to allow visual and olfactory habituation, in order to minimize neophagia prior to offering HFD for consumption. Mice were exposed to HFD pellets ad libitum for 85-125 minutes prior to perfusion, and the pellets were weighed before and after exposure to measure consumption. For HFD+FR stimuli, mice had 24 hours without food before being exposed to HFD pellets ad libitum for 85-125 minutes before perfusion. After 90 mins, mice were perfused and brains processed for c-Fos immunohistochemistry as described.

### Anterograde Tracing Analysis

To quantitatively assess anterograde outputs we performed confocal imaging of 4-6 slices spanning the PSTN and PVN (spaced at approximately 120 um intervals) and utilized open-source FIJI image analysis tools to acquire mean fluorescence values.

### Statistical Analyses

Food and alcohol intake values were analyzed by two-way repeated measures ANOVA (within-subjects factor of treatment [Saline, DCZ], between-subjects factor of Sex). Gq-DREADD cFos activation results were analyzed by two-way repeated measures ANOVA (within-subjects factor of treatment [Saline, DCZ], between-subjects factor of Virus). *Vipr2*-tomato cFos activation results were analyzed by two-way ANOVA (between-subjects factors of Group and Sex), absence of Sex effects justified presenting the analysis as one-way ANOVA, with multiple comparison-corrected Bonferroni post-hoc between-group comparisons. *Vipr2*-tomato PKCd immunostaining data were analyzed by two-way repeated measures ANOVA (within-subjects factor of Bregma level [Anterior, Middle, Posterior], between-subjects factor of Sex). *Vipr2* anterograde tracing data were analyzed by two-way repeated measures ANOVA (within-subjects factor of Bregma level [Anterior, Middle, Posterior], between-subjects factor of Sex). Analyses and graphs were generated using Graphpad Prism.

## RESULTS

### ovBNST^Vipr2^ chemogenetic activation promotes intake of food but not alcohol

To evaluate the role of ovBNST*^Vipr2^* neurons in consumption, we quantified intake of normal chow food and alcohol (20% EtOH) during daily consecutive 4hr sessions of two-bottle choice drinking in the circadian dark cycle (active phase) (**Fig. 1A**). Chemogenetic stimulation of ovBNST*^Vipr2^* neurons upon DCZ injection nearly doubled food intake in both sexes, relative to saline control sessions (**Fig. 1B**). However, ovBNST*^Vipr2^* stimulation had no significant effects on alcohol intake (**Fig. 1C**) or preference (**Fig. 1D**). To validate activation of ovBNST*^Vipr2^*neurons, mice received either saline or DCZ prior to perfusion. Using confocal imaging, we observed hM3Dq-mCherry and control virus expression limited to the oval subnucleus of the BNST **(Fig. S1A)**. Using cFos immunohistochemistry, we observed a significant increase in the proportion of DREADD expressing neurons co-expressing cFos, indicating specific activation of ovBNST*^Vipr2^* neurons by the Gq-coupled DREADDs **(Fig. S1B)**.

**Figure 1.**
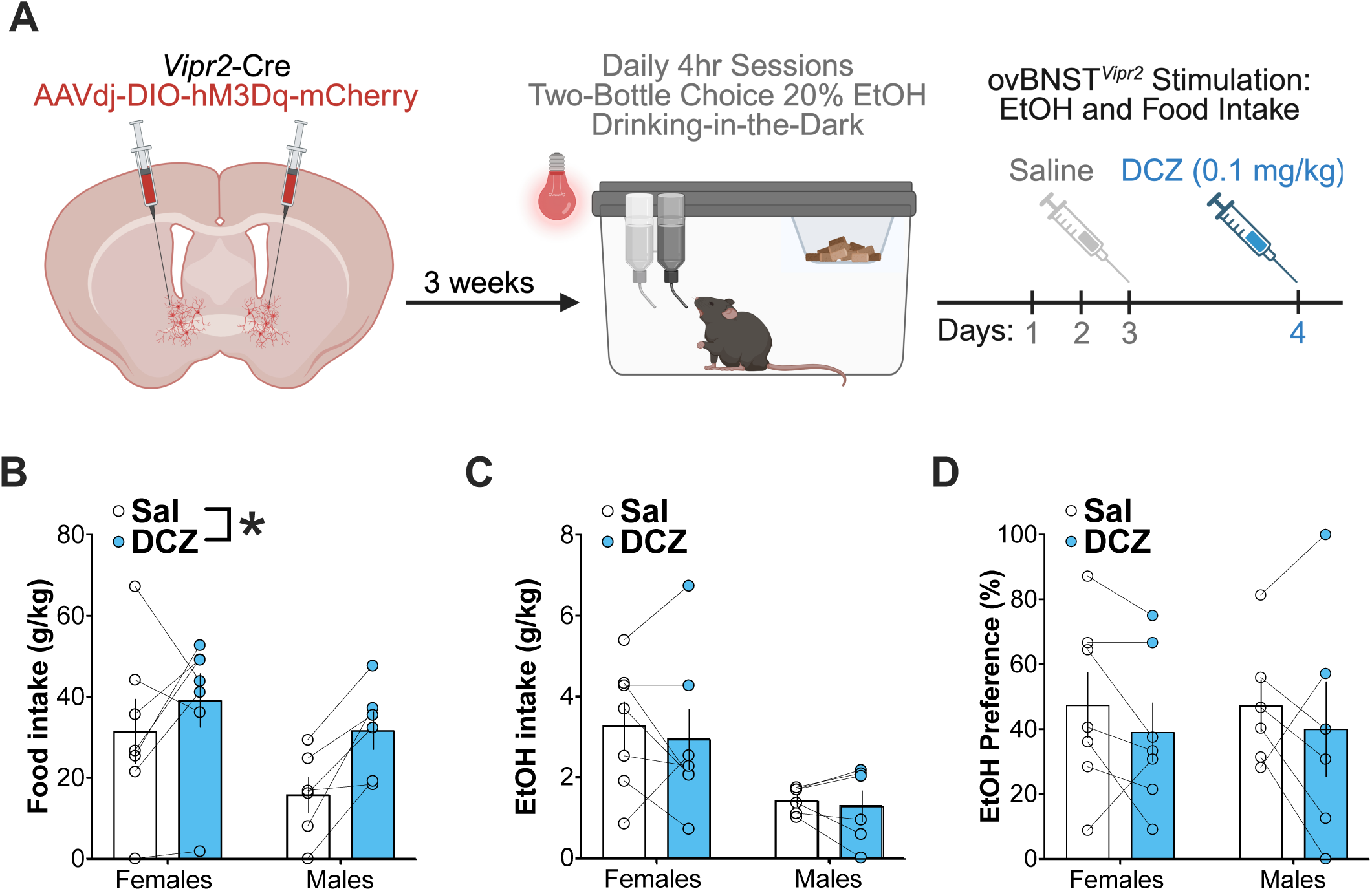
Chemogenetic activation of ovBNST-*Vipr2* neurons increases Food Intake. **A)** Schematic timeline of two-bottle choice DID experiment. Mice were given 3 weeks to recover from surgery and allow time for DREADD expression before beginning baseline drinking. For 4 consecutive days, mice were given access to an alcohol bottle and a water bottle for 4 hours early in the dark cycle. The following week, mice underwent the same procedure but received saline injections before each drinking session for the first 3 days and DCZ on the 4th day. **B)** Food intake normalized to body weight (two-way ANOVA; Main effect of DCZ; F_1,11_ = 7.962; *p* = 0.0166). **C)** Alcohol consumption normalized to body weight averaged across saline injection days and compared to the 4th day of DCZ injection (two-way ANOVA; F_1,11_ = 0.486; *p* = 0.499). **D)** Ethanol preference ratio defined by the volume of alcohol consumed divided by the total alcohol and water consumed (two-way ANOVA; F_1,11_ = 1.779; *p* = 0.2093).

### Vipr2 ovBNST neurons are strongly activated by food restriction

Upon observing increased food intake following ovBNST*^Vipr2^*DREADD stimulation, we aimed to determine how feeding-related stimuli impact ovBNST*^Vipr2^* neuronal activation patterns, using cFos immunohistochemistry in *Vipr2*-tomato fluorescent reporter mice. We examined cFos/Vipr2-tomato overlap in the ovBNST across 4 different groups of mice: CTL (control, no stimulus), high-fat diet consumption (HFD), food restriction (FR), and FR followed by HFD re-feeding (FR+HFD).

Relative to CTL conditions, we found that FR led to considerable cFos activation in ovBNST*^Vipr2^* neurons (**Fig. 2A-D**). Upon quantification, we confirmed that FR significantly activated ovBNST*^Vipr2^*neurons in a manner that was not reversed by acute HFD re-feeding, whereas HFD alone did not significantly impact ovBNST*^Vipr2^* neuronal activation (**Fig. 2E-F**). To confirm elevated HFD intake under FR, we documented significantly increased HFD intake in FR+HFD mice relative to HFD-only mice (**Fig. S2**).

**Figure 2.**
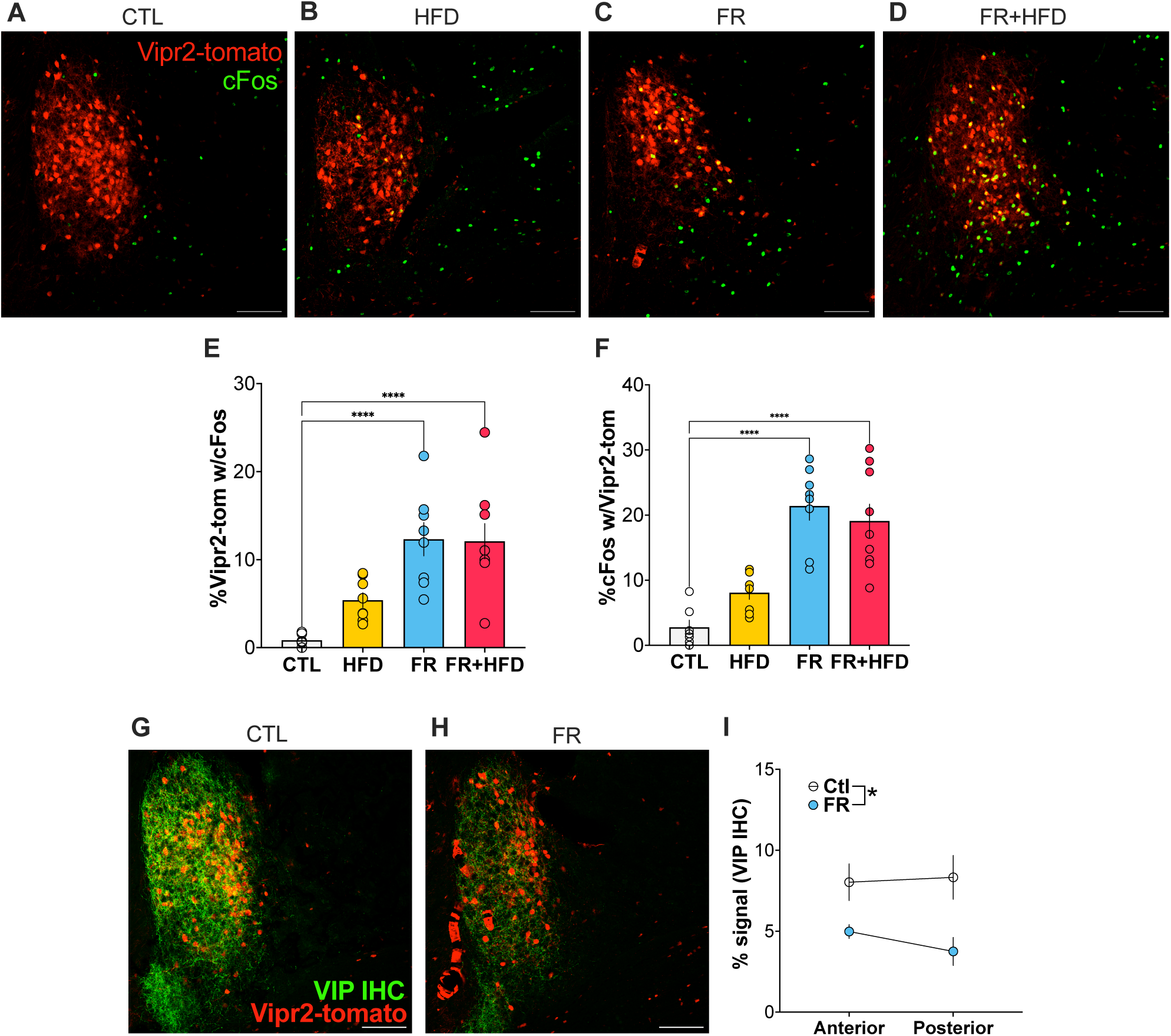
*Vipr2* ovBNST neurons are strongly activated by food restriction and display food restriction-induced changes in VIP innervation. **A-D)** Representative images of fluorescent immunohistochemistry (IHC) for cFos of various feeding-related stimuli. **E)** Quantification of %Vipr2-tomato neurons with cFos, demonstrating significantly increased ovBNST*^Vipr2^* cFos activation in FR and FR+HFD groups, relative to CTL (one-way ANOVA; F_3,28_ = 12.58, *p* <0.0001, Bonferroni post-hoc comparisons CTL vs. FR and CTL vs. FR+HFD both *p*<0.0001). **F)** Quantification of %cFos positive neuron overlap with Vipr2-tomato neurons demonstrating significantly increased ovBNST*^Vipr2^* cFos activation in FR and FR+HFD groups, relative to CTL (one-way ANOVA; F_3,28_= 20.00, *p* <0.0001, Bonferroni post-hoc comparisons CTL vs. FR and CTL vs. FR+HFD both *p*<0.0001)). **G-H)** Representative images of IHC for VIP in Ctl vs FR mice. **I)** Demonstration of significantly reduced VIP innervation in FR mice vs controls (F_1,5_ = 13.60, p=0.0142). Scale bars: 100um.

### Vipr2 ovBNST neurons display food restriction-induced changes in VIP innervation

Given that G-protein-coupled Vipr2 actions occur primarily via stimulatory Gs and Gq signaling, this suggests that increased VIP release and signaling may drive the observed ovBNST*^Vipr2^* activation during FR. To test this hypothesis, we performed immunohistochemistry (IHC) for VIP in ovBNST slices from CTL and FR mice, and we identified significantly decreased VIP axonal innervation of the ovBNST under FR conditions (**Fig. 2G-I**).

Since FR mice demonstrated increased ovBNST*^Vipr2^* cFos activation in conjunction with decreased VIP IHC signal, this suggested the possibility that decreased VIP signal was reflective of enhanced VIP release from presynaptic terminals. To explore this possibility, we performed correlational analyses of ovBNST*^Vipr2^* cFos activation levels with VIP IHC signals. Consistent with our hypothesis, we identified a significant negative correlation, in which greater ovBNST*^Vipr2^* cFos activation was associated with reduced VIP IHC signal (**Fig. S3**).

### Vipr2 and PKCd are largely non-overlapping ovBNST neuronal subpopulations

Given previous findings on ovBNST control of food intake, including reports in which stimulation of ovBNST neurons marked by protein kinase C delta (PKCd) inhibited feeding [9, 10], we set out to determine the potential overlap between *Vipr2* and PKCd neurons in the ovBNST. Fluorescent immunohistochemical staining for PKCd in *Vipr2*-tomato mice revealed that these markers label two largely distinct neuronal subpopulations in the ovBNST (10-15% overlap, **Fig. 3A-D**) The greatest degree of overlap was apparent only in posterior sections (25-35%). Furthermore, given reported sex differences in specific BNST subregions, neuronal subpopulations, and gene expression patterns [29–32], we directly compared cell counts and overlap profiles between female and male mice, finding no significant differences between sexes (**Fig. 3E-I**). Overall, these findings indicate that the opposing effects of ovBNST*^Vipr2^* and ovBNST*^PKCd^* stimulation on food intake (increasing and decreasing feeding, respectively), can be explained by their lack of overlap and formation of largely distinct neuronal subpopulations within the ovBNST.

**Figure 3.**
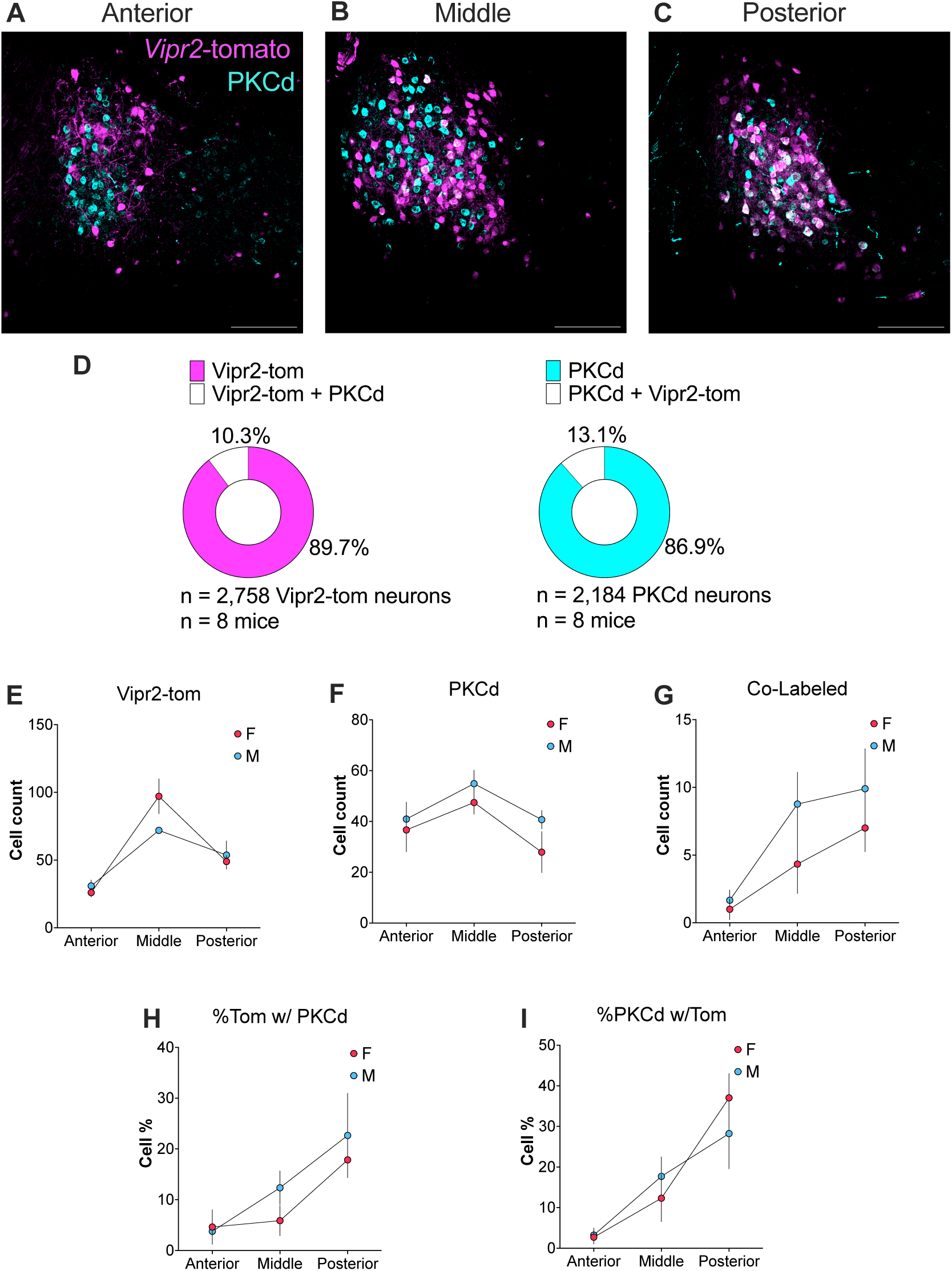
*Vipr2* and PKCd are largely non-overlapping ovBNST neuronal subpopulations. **A-C)** Representative images of fluorescent immunohistochemistry (IHC) for PKC-delta in the ovBNST of a *Vipr2*-tomato mouse across the anterior/posterior axis. Vipr2-tomato pseudocolored in Magenta, PKCd pseudocolored in Cyan, White = co-labeled neurons. Scale bar = 100um. **D)** Chart displaying quantification of Vipr2 and PKCd overlap (10-15%). **E-I)** Female and male mice did not significantly differ in cell counts for *Vipr2*-tomato, PKCd, or their overlap. Posterior ovBNST slices exhibited greater co-labeled *Vipr2*-tomato/PKCd neurons, relative to ovBNST slices from anterior and middle sections. Scale bars: 100um.

*Vipr2-BNST anterograde viral tracing identifies major outputs to the parasubthalamic nucleus (PTSN) and paraventricular nucleus of the hypothalamus (PVN)*.

To identify potential circuit mechanisms by which ovBNST*^Vipr2^*neurons promote feeding, we performed Cre-dependent anterograde viral tracing using the mGFP-2A-Synaptophysin mRuby virus (**Fig. 4A**), which fluorescently marks cells with two fluorophores that label either all cellular compartments (mGFP) or specifically presynaptic sites (mRuby). We identified primary axonal outputs from ovBNST*^Vipr2^*neurons to the parasubthalamic nucleus (PTSN) and paraventricular nucleus of the hypothalamus (PVN) (**Fig. 4B-D**), two structures that are both implicated in feeding. We also sought to identify potential sex differences in the strength of ovBNST*^Vipr2^* outputs, quantifying the signal strength in downstream regions of both female and male mice. We identified significantly greater innervation of the PVN (but not PSTN) in male vs. female mice, specifically at anterior levels, indicating that ovBNST*^Vipr2^* neurons may provide sex-specific signals to hypothalamic regions that regulate stress and feeding (**Fig. 4E-F**).

**Fig. 4.**
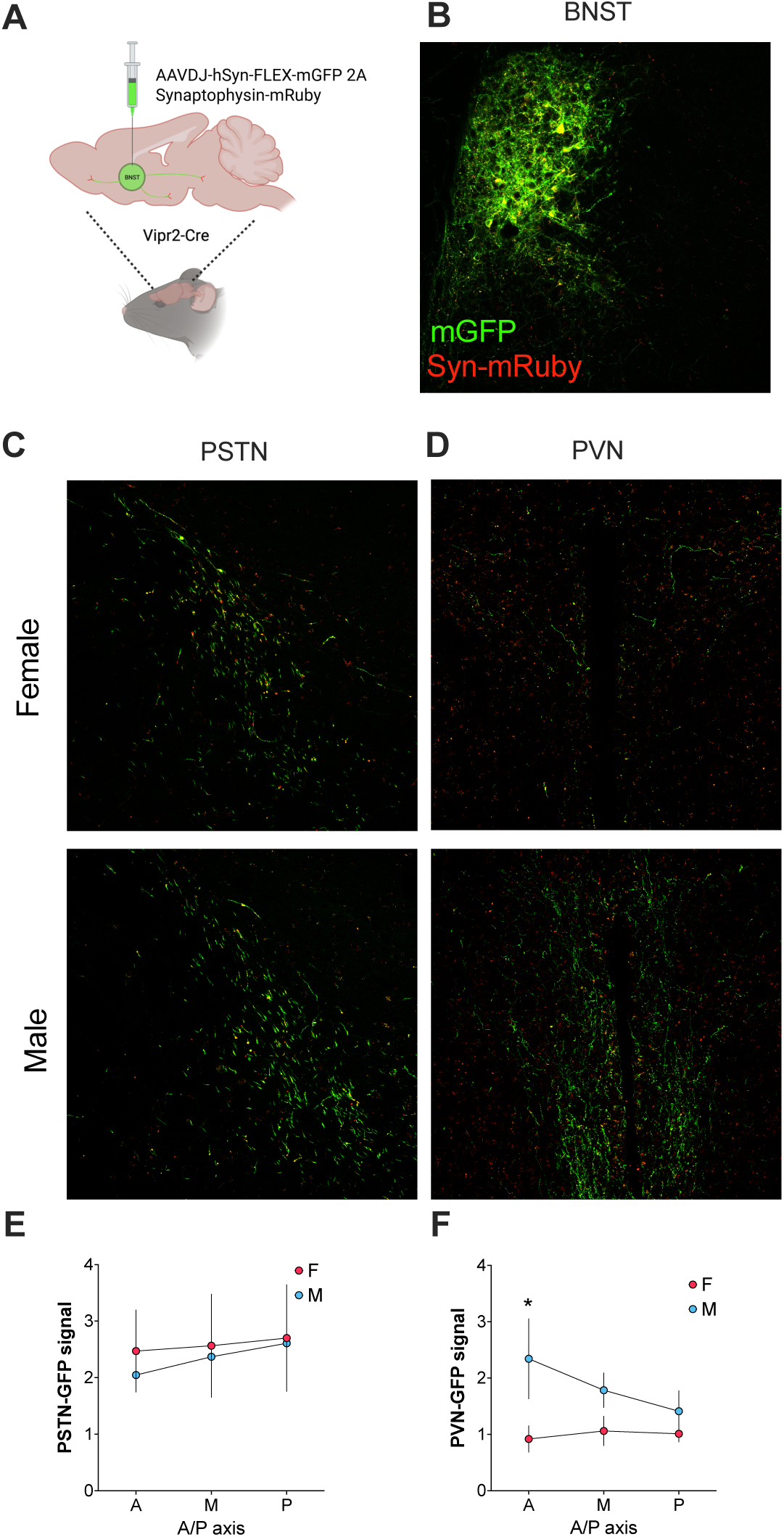
V*i*pr2-BNST anterograde viral tracing identifies major outputs to the parasubthalamic nucleus (PTSN) and paraventricular nucleus of the hypothalamus (PVN). **A)** Schematic of viral Synaptophysin-mRuby surgeries **B)** Representative images of mGFP and mRuby fluorescence in ovBNST*^Vipr2^* neuronal cell bodies. **C-D)** Representative images of mGFP and mRuby fluorescence in the PSTN and PVN of females and males. **E-F)** Quantification of mGFP fluorescent signals in the PSTN and PVN of females and males, demonstrating increased mGFP signals within the PVN of male mice in the anterior regions (Sex x Bregma interaction; F_2,14_ = 3.113, *p* = 0.0761, \**p*=0.0289 anterior Female vs. Male). Scale bars: 100um.

## DISCUSSION

In this study, we characterized the population of Vipr2-expressing neurons within the ovBNST and our results indicate that ovBNST*^Vipr2^*neurons promote food intake and are strongly modulated by food availability. We demonstrated that selective chemogenetic activation of ovBNST*^Vipr2^*significantly increased chow consumption, and that food restriction (FR) significantly increased cFos co-expression in ovBNST*^Vipr2^* neurons, indicative of robust neuronal activation by homeostatic challenge. Complementary IHC experiments revealed that FR also decreased axonal VIP immunoreactivity in the ovBNST in a manner that was negatively correlated with FR-induced ovBNST*^Vipr2^* cFos expression, suggestive of FR-induced VIP release and stimulatory Vipr2 signaling in the ovBNST. Finally, we showed that ovBNST*^Vipr2^* neurons are largely non-overlapping with a previously characterized ovBNST subpopulation marked by protein kinase C delta (PKCδ/*Pkcd*), and that they display prominent axonal projections to the PVN and PSTN of the hypothalamus, with some evidence for sex differences in their projection patterns. These findings suggest that ovBNST^Vipr2^ neurons represent a distinct feeding-promoting subpopulation within the extended amygdala that is dynamically regulated by both metabolic cues and neuropeptidergic signaling.

Our results add to a body of evidence supporting the idea that the BNST contains distinct cell types with divergent influences on feeding. Prior work found that stimulation of ovBNST*^Prkcd^*neurons suppressed feeding [9, 10], in opposition to the orexigenic effects of stimulating the broader BNST*^Vgat^* population that projects to the lateral hypothalamus [5, 7, 8]. We found that Vipr2 and PKCδ neurons largely constitute non-overlapping subpopulations, providing a cellular basis for these opposing functions within the same anatomical subnucleus. This functional counterbalance resembles other extended amygdala and hypothalamic circuits in which parallel excitatory and inhibitory pathways regulate motivated behaviors in opposing directions [12, 25, 33]. By delineating the distinct contributions of these ovBNST subpopulations, our findings highlight the importance of cellular heterogeneity in shaping complex behaviors such as feeding.

### VIP/Vipr2 Signaling in Feeding and Appetite

Given our finding that VIP innervation is decreased under food restriction, one mechanism is that ovBNST*^Vipr2^* neurons are situated in a broader VIP circuit. Global genetic knockout of VIP led to reduced body weight attributed to disrupted circadian rhythms in feeding, but the neural substrates underlying VIP’s effects on feeding were not investigated [34]. Analogously, VIP is generated in several neuronal populations (e.g., cortex, suprachiasmatic nucleus, midbrain), but VIP signaling in the BNST has not been directly studied as a neuromodulatory system in feeding behavior. The source of VIP input to the BNST is likely from the PAG [35], but previous studies that explored the PAG➜BNST pathway focused on the dopaminergic neurons in pain processing [36]. Regarding the source of VIP into the ovBNST, it remains unclear whether our observed FR-induced decrease in axonal VIP immunoreactivity is due to decreased VIP synthesis at the cell body level, or a depletion in axon terminals resulting from extensive VIP release. Consistent with stimulatory VIP-Vipr2 signaling through canonical Gs-or Gq-coupled pathways, FR-induced ovBNST*^Vipr2^* cFos activation is suggestive of FR-induced axonal VIP release. Further work is required to understand how food restriction and metabolic changes modulate VIP release and signaling in the ovBNST.

### BNST-Hypothalamic Circuitry in Feeding

Anatomical anterograde tracing revealed that neurons ovBNST*^Vipr2^*neurons project prominently to both the parasubthalamic nucleus (PSTN) and the paraventricular nucleus of the hypothalamus (PVN), two regions with well-established roles in feeding. Prior work found that activation of the BNST➜PVN pathway increased rewarding seeking behavior [37], consistent with the idea that food is especially rewarding in the fasted state. Various cell types in the PSTN mediate appetite suppression [38] and delayed refeeding [39]. Kim et al. specifically showed that activation of PSTN*^Tac1^* neurons decreased food intake in fasted mice [38]. Given that the BNST is primarily composed of GABAergic neurons, it remains possibly that ovBNSTVipr2 neurons can promote food consumption by inhibition PSTN Tac1 neurons.

### Clinical Interpretations of Stress Circuits and Feeding

The neurobiological mechanisms governing feeding in the context of psychiatric disorders have been of significant interest to researchers and clinicians. Stress, both acute and chronic, has been known to contribute to disordered eating. Interestingly, the effect of stress on eating has significant heterogeneity, with a roughly equal number of people eating more or less in response to stress [40]. Thus, there is a critical need to understand the mechanisms underlying individual variation in eating behavior following stress. For example, one speculative hypothesis is that ovBNST*^Vipr2^* stimulation could be a viable therapeutic strategy to overcome stress-related anorexia. Indeed, recent work discovered that the appetite-reducing weight loss compound Semaglutide (Glp1r agonist) strongly activated ovBNST*^Prkcd^* neurons [5], suggesting that ovBNST^Vipr2^ neuron activation may present a target for counteracting GLP1 actions in the ovBNST and minimizing problematic weight loss in conditions of cachexia or anorexia.

The BNST is enriched in steroid hormone receptors and demonstrates sexual dimorphism at the level of multiple unique neuropeptide subpopulations [30–32, 41–43], suggesting the BNST may also display sex-specific mechanisms contributing to food consumption. Thus, we analyzed data from both female and male mice in terms of ovBNST*^Vipr2^* effects on feeding, activity modulation by food restriction, and outputs to hypothalamic structures.

### Limitations

We note that our study has some limitations. First, while projections to the PVN and PSTN were identified, the functional relevance of these ovBNST*^Vipr2^* pathways in feeding or refeeding remains largely unexplored; targeted manipulations will be necessary to determine whether these connections are necessary or sufficient for the observed feeding effects. Second, the limited numbers of female and male subjects may obscure potentially greater sex differences in circuit organization and function, and studies of VIP innervation based on feeding behavior were performed only in female mice. Third, the precise time course of DCZ-induced feeding as well as FR-induced changes in ovBNST*^Vipr2^* VIP innervation remain unknown and require further investigation to temporally link these processes. Additionally, all chemogenetic behavioral data were collected during concurrent alcohol availability (though chemogenetic manipulation of ovBNST*^Vipr2^* neurons did not produce significant changes in alcohol drinking). Finally, VIP immunohistochemistry provides only indirect evidence of peptide release; direct measurement of VIP release and signaling dynamics using pharmacology, fiber photometry, or related techniques would strengthen the interpretation of these findings.

In conclusion, our results identify ovBNST*^Vipr2^* neurons as a previously uncharacterized BNST subpopulation that promotes food intake, is activated by feeding restriction, and links BNST neuropeptide signalling to hypothalamic feeding circuits. Future research dissecting the projection specific roles and sex-dependent differences of this circuit will further illuminate its contributions to adaptive and maladaptive feeding.

## ACKNOWLEDGEMENTS

This work was supported by funding from NIH/NIAAA R00AA025677 (W.J.G.), Whitehall Foundation, Brain Research Foundation (W.J.G.). We thank Haniyyah Sardar for contributions to initial experiments and all Giardino Lab members for their support and assistance.

## CONFLICT OF INTEREST

Authors declare there are no competing financial interests in relation to the work described.

## Notes

### Competing Interest Statement

The authors have declared no competing interest.

