## Supplemental Figures for "A Distinct Subpopulation of Extended Amygdala Neurons Drives Food Intake"

Short Title: Vipr2 ovBNST Neurons Promote Feeding

Isaac F. Kandil[1]*, Ethan T. Rogers[1]*, Allison R. Morningstar[1], William J. Giardino[1‡]

[1]Wu Tsai Neurosciences Institute and Department of Psychiatry and Behavioral Sciences, Stanford University School of Medicine

*I.F.K. and E.T.R. contributed equally

‡Correspondence should be addressed to:

Will Giardino
Department of Psychiatry and Behavioral Sciences
3165 Porter Dr.
Palo Alto, CA 94304


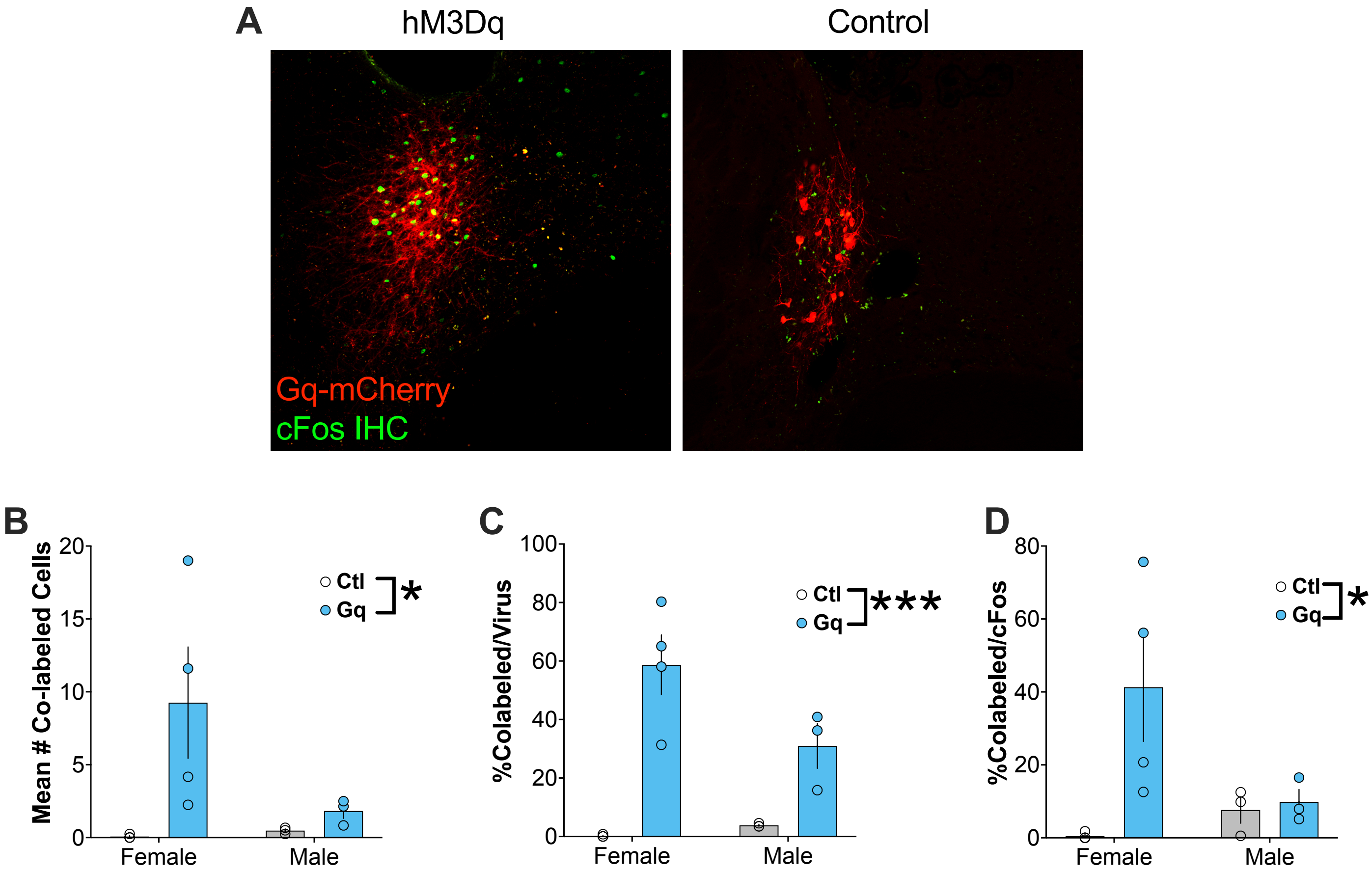


**Figure S1.** ***Vipr2-BNST*** **DREADD validation**

**A-B)** Representative images of mice expressing mCherry and IHC for cFos in the ovBNST of *Vipr2*-Cre mice that received either hM3Dq tagged with mCherry or mCherry alone. **C)** Mean number of cells expressing AAV-mCherry and co-labeled with cFos per mouse (two-way ANOVA with *main effect of hM3Dq virus; $F_{1,10} = 5.380$, $p = 0.0428$). **D)** Average percentage of colabeled cells normalized to the number of virus expressing cells (two-way ANOVA with ***main effect of hM3Dq virus; $F_{1,10} = 39.06$, $p < 0.0001$). **E)** average percentage of colabeled cells normalized to the number of cFos expressing cells (two-way ANOVA with *main effect of hM3Dq virus; $F_{1,10} = 5.673$, $p = 0.0385$).

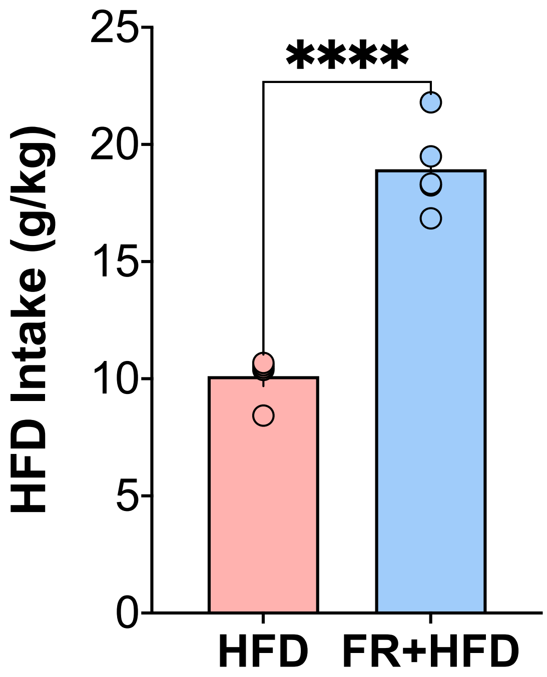


**Figure S2. High Fat Diet intake**

Comparison of high fat diet intake between groups which received HFD after no food restriction vs food restricted mice, demonstrating significantly increased consumption in food restricted mice (Welch's two-tailed t-test, $t=9.507$, $df=5.953$, ****$p < 0.0001$).

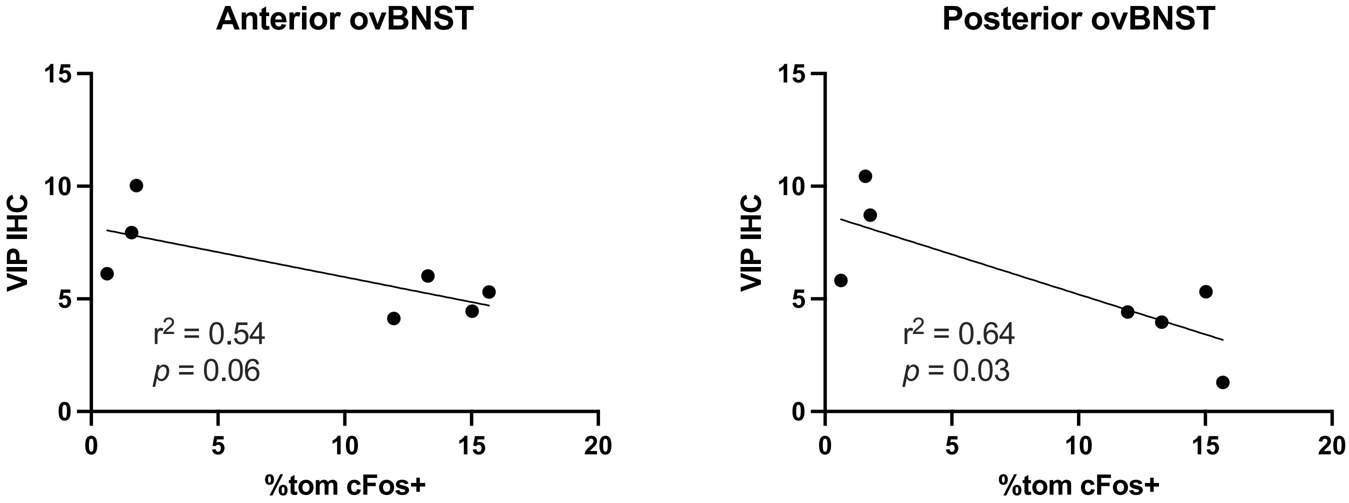


**Figure S3. *Vipr2*-tomato/VIP IHC Correlation**

Correlation between ovBNST VIP IHC signal and %Vipr2-tom cFos+ neurons, demonstrating a significant negative correlation in Posterior BNST sections ($F_{1,5} = 8.951$, $p = 0.0304$, $r^2 = 0.6416$).
